# Disruption of cortical dopaminergic modulation impairs preparatory activity and delays licking initiation

**DOI:** 10.1101/337014

**Authors:** Ke Chen, Roberto Vincis, Alfredo Fontanini

## Abstract

Dysfunction of motor cortices is thought to contribute to motor disorders such as Parkinson’s disease (PD). However, little is known on the link between cortical dopaminergic loss, abnormalities in motor cortex neural activity and motor deficits. We address the role of dopamine in modulating motor cortical activity by focusing on the anterior lateral motor cortex (ALM) of mice performing a cued-licking task. We first demonstrate licking deficits and concurrent alterations of spiking activity in ALM of mice with unilateral depletion of dopaminergic neurons (i.e., mice injected with 6-OHDA into the medial forebrain bundle). Hemi-lesioned mice displayed delayed licking initiation, shorter duration of licking bouts, and lateral deviation of tongue protrusions. In parallel with these motor deficits, we observed a reduction in the prevalence of cue responsive neurons and altered preparatory activity. Acute and local blockade of D1 receptors in ALM recapitulated some of the key behavioral and neural deficits observed in hemi-lesioned mice. Altogether, our data show a direct relationship between cortical D1 receptor modulation, cue-evoked and preparatory activity in ALM, and licking initiation.

**SIGNIFICANCE STATEMENT:** The link between dopaminergic signaling, motor cortical activity and motor deficits is not fully understood. This manuscript describes alterations in neural activity of the anterior lateral motor cortex (ALM) that correlate with licking deficits in mice with unilateral dopamine depletion or with intra-ALM infusion of dopamine antagonist. The findings emphasize the importance of cortical dopaminergic modulation in motor initiation. These results will appeal not only to researchers interested in cortical control of licking, but also to a broader audience interested in motor control and dopaminergic modulation in physiological and pathological conditions. Specifically, our data suggest that dopamine deficiency in motor cortex could play a role in the pathogenesis of the motor symptoms of Parkinson’s disease.

## INTRODUCTION

Dysfunction of motor cortices, which are important for movement planning, initiation and execution, has been suggested to play a role in the motor symptoms of Parkinson’s disease (PD) (Lindenbach and Bishop, 2013). Studies on motor cortices of human patients and animal models of PD revealed abnormalities in preparatory activity, excitability, excitation/inhibition balance and oscillatory dynamics (Doudet et al., 1990; Ridding et al., 1995; Goldberg et al., 2002; Escola et al., 2003; Lefaucheur, 2005; Pasquereau and Turner, 2011; Pasquereau et al., 2015). However, it is unclear whether abnormal patterns of motor cortical activity are secondary to dysfunction of the basal ganglia or whether they result from disruption of local dopaminergic modulation. Midbrain dopaminergic neurons project to the striatum and motor cortex. While dopaminergic innervation to the striatum has been studied extensively for its modulatory role on motor initiation and execution, studies on dopaminergic innervation to the motor cortex have been more limited and focused mostly in its role in synaptic plasticity and motor skill learning (Molina-Luna et al., 2009; Guo et al., 2015). To date, little is known about the direct link between loss of dopaminergic signaling in the motor cortex, alterations of motor cortical single unit activity, and corresponding motor deficits.

Here, we investigate the role of motor cortex dopaminergic transmission in movement initiation and execution. We focus on the anterior lateral motor cortex (ALM) of mice engaged in a cued-licking task. Licking was chosen because it is an innate motor behavior whose cortical control is well-studied. In rodents, licking is regulated by a central pattern generator circuit in the brainstem, which is under the control of the motor cortex (Travers et al., 1997). ALM plays an important role in the planning and execution of licking (Komiyama et al., 2010; Guo et al., 2014; Li et al., 2015; Inagaki et al., 2018), as reflected by the presence of neurons whose firing rates are modulated before the onset of licking (defined as “preparatory” neurons) (Guo et al., 2014; Li et al., 2015; Chen et al., 2017; Inagaki et al., 2018). In addition, this area appears to be responsible for controlling the direction of tongue movements, as unilateral optogenetic silencing of ALM can introduce a directional bias towards the ipsilateral side (Guo et al., 2014; Li et al., 2015). Although ALM has been studied for its involvement in controlling normal licking, how lack of dopaminergic signaling impacts activity and function of this region remains unknown.

The experiments described here rely on behavioral training, pharmacology, and electrophysiological recordings to study licking deficits and related abnormalities of ALM neural activity in the context of unilateral dopamine depletion (i.e., unilateral injection of 6-OHDA into the medial forebrain bundle). This manipulation has been classically used to model some of the features of PD (Lundblad et al., 2004; Thiele et al., 2012; Jagmag et al., 2016). First, we show that mice with unilateral dopamine depletion display delayed licking initiation, shorter duration of licking bouts, and deviated tongue protrusions compared to control mice. Next, we report changes in cue responses and preparatory activity for neurons in ALM of 6-OHDA lesioned mice. Finally, we perform local pharmacological blockade of dopaminergic receptors to determine the contribution of cortical dopaminergic deficit in ALM to the electrophysiological and behavioral alterations seen in 6-OHDA lesioned mice.

Using licking as a model behavior, our data show motor deficits and abnormalities in neural activity associated with unilateral dopamine depletion. The results demonstrate the importance of cortical dopaminergic modulation for motor initiation and for modulating preparatory activity.

## MATERIALS AND METHODS

### Experimental subjects

The experiments were performed on adult male mice (C57BL/6, 12-20 weeks old, Charles River). Mice were group housed and maintained on a 12 h light/dark cycle with *ad libitum* access to food and water unless otherwise specified. All experimental protocols were approved by the Institutional Animal Care and Use Committee at Stony Brook University, and complied with university, state, and federal regulations on the care and use of laboratory animals.

### Surgical procedures for 6-OHDA injections in the medial forebrain bundle

Mice were anesthetized with isoflurane (1-1.5%) in oxygen (1 L/min). Once fully anesthetized, mice were placed on a stereotaxic apparatus. The scalp was cut open to expose the skull and a hole was drilled above the medial forebrain bundle (MFB, anterior-posterior: −1.2 mm, medial-lateral: 1.3 mm, dorsal-ventral: −4.75 mm). In a first group of mice (referred hereafter as 6-OHDA lesioned), 3.5 μg 6-OHDA dissolved in 1 μl 0.02% ascorbic acid (vehicle, prepared from sterile saline) was unilaterally injected into the MFB. A second group of mice (sham-lesioned mice, referred hereafter as control) underwent the same surgical procedure but received 1 μl vehicle injection into the MFB. To prevent dehydration, mice were monitored daily and subcutaneously injected with 1 mL lactated ringer’s solution after the surgery as needed. In addition, food pellets soaked in 15% sucrose were placed on the floor of cages to facilitate eating (Francardo et al., 2011).

### Behavioral screening of lesion: cylinder test

Two to three weeks after the MFB lesion surgery, mice were placed into a clear plastic cylinder. Mice could freely explore the cylinder, rearing and touching the cylinder wall with their forepaws. The behavior during the first 3 min in the cylinder was videotaped and analyzed. The number of wall touches with the ipsilateral or contralateral forepaw was counted and used to calculate the forepaw preference. Only lesioned mice with less than 40% usage of contralateral forepaw for touching the cylinder wall were used for further experiments (Lundblad et al., 2004).

### Surgical procedures for implanting electrodes, infusion cannula, and electrode-cannula assemblies

2-4 weeks after the lesion surgery, 6-OHDA lesioned and control mice were anesthetized with an intraperitoneal injection of a mixture of ketamine (70 mg/kg) and dexmedetomidine (1 mg/kg) and placed on a stereotaxic apparatus. The scalp was incised to expose the skull. For electrode implantation, 1 mm craniotomies were performed above both anterior lateral motor cortices (ALM, anterior-posterior: 2.4 mm, medial-lateral: ±1.5 mm) and two holes were drilled above visual cortex on both hemispheres for inserting ground wires (silver wire). A linear array of 16 electrodes (formvar-insulated nichrome wire, catalog no. 761000, A-M System, Sequim, WA) was bilaterally implanted into ALM (dorsal-ventral: −0.8 ‑ −1 mm). For infusion cannula implantation, naïve mice were used instead, and a 1 mm craniotomy was performed on left ALM. A 26-gauge guide cannula with a dummy (0.5 mm projection) was inserted into ALM (dorso-ventral: −700 μm). To record single units after local D1 receptor blockade, a group of naïve mice was unilaterally implanted in ALM with a custom-built ensemble containing 8 tetrodes (Item No. PX000004, Sandvik-Kanthal, Hallstahammar, Sweden) around an infusion guide cannula (26 gauge). Electrodes, cannulae or electrode-cannula assemblies and a head bolt (for the purpose of head restraint) were cemented to the skull with dental acrylic. Mice were allowed to recover from surgery for a week before starting water restriction regimen.

### Cued-licking paradigm

Following recovery, mice were started on a water restriction regime, with 1.5 ml water daily one week before training. Weight was monitored and maintained at > 80% of the standard weight for age, strain and sex. In the first phase of training, mice were habituated to restraint. During brief restraint sessions, a spout containing a drop of sucrose (200 mM) was moved close to the animal to encourage licking. Once the mouse started to reliably lick the spout, session duration was increased and training in the cued-licking paradigm began. For each trial, a movable spout containing a drop of sucrose (~3 μl, 200 mM) moved in front of the mouth of the animal 1 s after the onset of an auditory cue (200 ms, 2k Hz, 70 dB). The spout remained in place for 2 s to allow the mouse to lick and access the sucrose solution before retracting. The inter-trial interval was 10 s. An infrared beam (940 nm, powered by a fiber-coupled LED, Thorlabs, Newton, NJ) was put in front of the mouth of the mouse such that each lick could be detected. Orofacial movements were also recorded with a videocamera (30 Hz frame rate) synchronized with the data acquisition software (CinePlex, Plexon, Dallas, TX).

### Electrophysiological recordings in control and 6-OHDA lesioned mice

Multiple single units were recorded via a multichannel acquisition processor (Plexon) in mice performing the cued-licking paradigm. Neural signals were amplified, bandpass (300-8000 Hz) filtered, and digitized at 40k Hz. Single units were isolated by threshold detection and a waveform matching algorithm and were further sorted offline through principal component analysis using Offline Sorter (Plexon).

### D1/D2 receptor antagonist infusion in ALM

Thirty to forty minutes before a testing session, mice previously trained in the cued-licking paradigm were briefly anesthetized with 1% isoflurane and a 33-gauge inner cannula (0.5 mm projection) was inserted into the guide cannula. 0.5 μl of a solution of either the D1 receptor antagonist (5 μg/μl SCH23390 hydrocloride, Sigma-Aldrich, St. Louis, MO), the D2 antagonist (5 μg/μl raclopride tartrate salt, Sigma-Aldrich) or sterile saline (0.9%) was unilaterally infused into ALM at 0.25 μl/min using a syringe pump (11 plus, Harvard Apparatus, Holliston, MA).

### D1 receptor antagonist infusion in ALM and electrophysiological recordings

After recovery from the surgery for at least a week, mice were water restricted and trained to perform the cued-licking paradigm. Testing started after 8-12 days of training. Thirty to forty minutes before a testing and electrophysiological recording session, mice were head restrained and a 33-gauge inner cannula (0.5 mm projection) was inserted into the guide cannula. 0.5 μl of a solution of either the D1 receptor antagonist (5 μg/μl SCH23390 hydrocloride, Sigma-Aldrich) or sterile saline (0.9%) were infused into ALM at 0.25 μl/min using a syringe pump (11 plus, Harvard Apparatus). Single units were recorded and sorted offline as described above. Each session of saline infusion was followed, on the day after, by a session with D1 receptor antagonist infusion. Each mouse underwent 1-2 sessions of saline and SCH23390 infusion.

### Data analysis

Data analysis was performed using Neuroexplorer (Plexon) and custom written scripts in MATLAB (MathWorks, Natick, MA).

#### Analysis of licking behavior

The analog trace from the infrared beam (and its breaking by the tongue) was used for analyzing licking behaviors. A licking event was detected whenever the trace crossed a fixed threshold. A bout was defined as a train of at least three consecutive licks with an inter-lick interval shorter than 500 ms (Davis and Smith, 1992). Only licking bouts within 4 s after the auditory cue were used for the analysis. In the case of two licking bouts occurred in the same trial, only the first licking bout was used for analysis. Video analysis of the oral region was used to extract the angle of tongue protrusions at each lick. Licking angle was defined as an angle between the midline of the protruded tongue and the midline of the mouse chin.

#### Analysis of single unit

Single unit spike timestamps were aligned to either the onset of the auditory cue or the licking bout initiation. Perievent rasters of individual units were used to construct peristimulus time histograms (PSTHs, bin size is 100 ms). For analyzing population PSTHs, the firing rate of each neuron was normalized using area under the receiver operating characteristic curve (auROC) method (Cohen et al., 2012; Gardner and Fontanini, 2014). This method normalizes firing rate to a value between 0 and 1, in which 0.5 represents baseline firing rate, value > 0.5 or < 0.5 represents increased or decreased firing rate compared to the baseline, respectively. Population PSTH was calculated by averaging auROC across each unit.

#### Analysis of cue response

PSTHs of single units were aligned to onset of cue. Activity after onset of cue was assessed by examining firing activity in a 500 ms window after cue onset. Firing rates within each bin (bin size is 100 ms) in the 500 ms window after cue onset were compared to baseline (1 s before the auditory cue) with a Wilcoxon rank sum test (p < 0.05) and a correction for multiple comparison (Šidák correction).

#### Analysis of preparatory response

PSTHs of single units were aligned to bout initiation. Activity preceding licking (i.e., preparatory activity) was assessed by examining firing rates in a 500 ms window before bout initiation. Firing rates within each bin (bin size is 100 ms) in the 500 ms window before bout initiation were compared to baseline (1 s before the auditory cue) with a Wilcoxon rank sum test (p < 0.05) and a correction for multiple comparison (Sidak correction). Units with significantly increased firing rate before bout initiation were defined as “excitatory preparatory” units, where units with significantly decreased firing rate before bout initiation were deemed as “inhibitory preparatory”. The latency of preparatory activity of each neuron was computed based on “change point” (CP) analysis (Jezzini et al., 2013; Liu and Fontanini, 2015; Vincis and Fontanini, 2016). To calculate latency of preparatory activity relative to the cue or bout initiation, we aligned spikes to cue onset or bout initiation and computed the cumulative distribution (CDF) of spike occurrence across all trials in the time interval starting 2 s before and ending 4 s after the cue or bout initiation, respectively. A sudden change of firing rate caused a correspondent change of the slope of CDF and the occurrence of a CP. The timing of the first significant CP was defined as the latency of preparatory activity. For analysis of latency relative the cue onset, neurons without CP (8/307) or neurons with first CP (2/307) occurring later than 3s after the cue were excluded for the analysis. For analysis of latency relative to the licking initiation, neurons without CP (6/307) or neurons with first CP (11/307) occurring after the licking initiation were excluded.

### Histological staining for verification of lesions and electrode/canula positioning

Mice were deeply anesthetized with an intraperitoneal injection of a mixture of ketamine/dexmedetomidine at 2-3 times the anesthetic dose and were intracardially perfused with PBS followed by 4% paraformaldehyde. The brain was further fixed with 4% paraformaldehyde overnight and cryoprotected with 30% sucrose for 3 days. The brain was eventually cut with a cryostat into 50 or 80 coronal slices. For visualizing electrode and canula tracks, 80 slices were stained with Hoechst 33342 (1:5000 dilution, H3570, ThermoFisher, Waltham, MA) using standard techniques. For immunostaining of tyrosine hydroxylase, 50 μm slices were first incubated for 1 h with blocking solution (a mixture of 5% BSA, 5% normal goat serum and 0.02% Triton-X in PBS) and were then incubated overnight at 4 °C with primary antibody (rabbit anti-tyrosine hydroxylase, 1:1000 dilution, ab112, abcam, Cambridge, United Kingdom). Slices were washed with PBS, incubated for 4h at 4 °C with secondary antibody (Alexa Fluor 594 goat anti-rabbit IgG, 1:500 dilution, R37117, ThermoFisher), and finally stained with Hoechst 33342.

## RESULTS

We unilaterally injected 6-hydroxydopamine (6-OHDA) into the medial forebrain bundle (MFB) of mice to deplete dopaminergic neurons. 6-OHDA causes a unilateral depletion of dopaminergic fibers in the striatum and loss of dopaminergic neurons in ventral tegmental area (VTA) and substantia nigra pars compacta (SNc) (**Figure 1A** and **1B**) (Lundblad et al., 2004; Thiele et al., 2012). The effectiveness of the lesion was assessed by comparing the number of weight bearing wall touches between the ipsilateral and contralateral forelimbs with a cylinder test (**Figure 1C**) (Schallert et al., 2000; Lundblad et al., 2002). Lesioned mice show a lower percentage of touches with the contralateral forelimb compared to intact mice (Lundblad et al., 2004). In accordance with the literature (Lundblad et al., 2004; Lundblad et al., 2005), we screened mice with motor deficits and included them in the study only if they showed less than 40% usage of the contralateral paw compared to control (**Figure 1D**). We confirmed the loss of dopaminergic neurons and fibers with histological staining.

**Figure 1.**
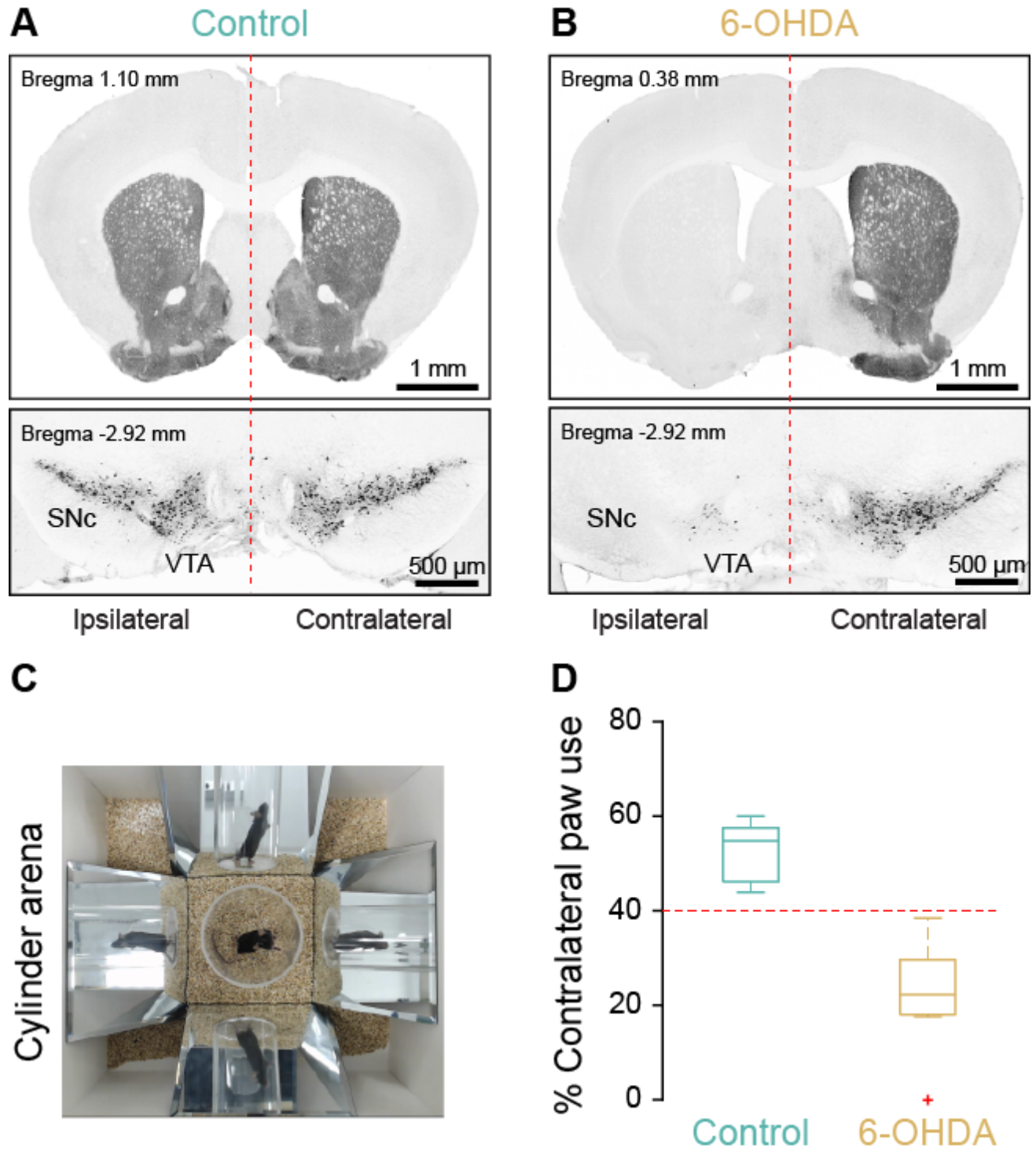
Confirmation of lesion and motor deficits after unilateral 6-OHDA injections in MFB. **A and B**, Representative tyrosine hydroxylase (TH) immunofluorescence staining showing dopaminergic fibers in striatum (top panel) and dopaminergic neurons in SNc and VTA (bottom panel) in a control mouse (**A**) and in a 6-OHDA lesioned mouse (**B**). Vertical dashed red lines indicate the midline of the brain. **C**, A representative snapshot of a unilateral 6-OHDA lesioned mouse performing the cylinder test. **D**, Boxplots of percentage of contralateral paw usage during the cylinder test in control (n = 9, blue) and screened 6-OHDA lesioned mice (n = 7, brown).

### Licking deficits with dopamine depletion

To assess for possible deficits in licking behaviors, 6-OHDA lesioned mice (n = 7) and vehicle injected control mice (n = 9) were trained to lick a spout to receive a drop of sucrose 1 s after an anticipatory auditory cue (**Figure 2A**). **Figure 2B** and **2C** show raster plots of licks from control and 6-OHDA lesioned mice, respectively. We analyzed the latency and duration of licking bouts (**Figure 2D**). The latency of bout initiation was significantly longer in lesioned mice compared to controls (2.40 ± 0.08 s vs 1.06 ± 0.04 s, *t*_(14)_ = 15.78, *p* = 2.6 × 10^−10^) (**Figure 2E**). The bout duration was shorter in lesioned mice relative to controls (1.05 ± 0.06 s vs 1.70 ± 0.062 s, *t*_(14)_ = −7.24, *p* = 4.3 × 10^−6^) (**Figure 2F**). The inter-lick interval, however, was not significantly affected (6-OHDA lesion vs control: 138.8 ± 4.1 ms vs 144.8 ± 4.5 ms, *t*_(14)_ = −1, *p* = 0.336). In addition to the timing, we also assessed the direction of tongue movements during licking via analysis of videos of the orofacial region (**Figure 2G**). The direction of tongue movements was quantified by calculating the angle between the axis of symmetry of the tongue and the midline of the mouth (see methods). A positive angle indicated a directional bias toward the side ipsilateral to the lesion, whereas a negative angle indicated a contralateral bias. 6-OHDA lesioned mice showed a positive licking angle that was significantly different from that observed in control mice (27.9 ± 5.8 deg vs −0.6 ± 1.0 deg, Welch’s t-test, *t*_(6)_ = −4.82, *p* = 0.003).

**Figure 2.**
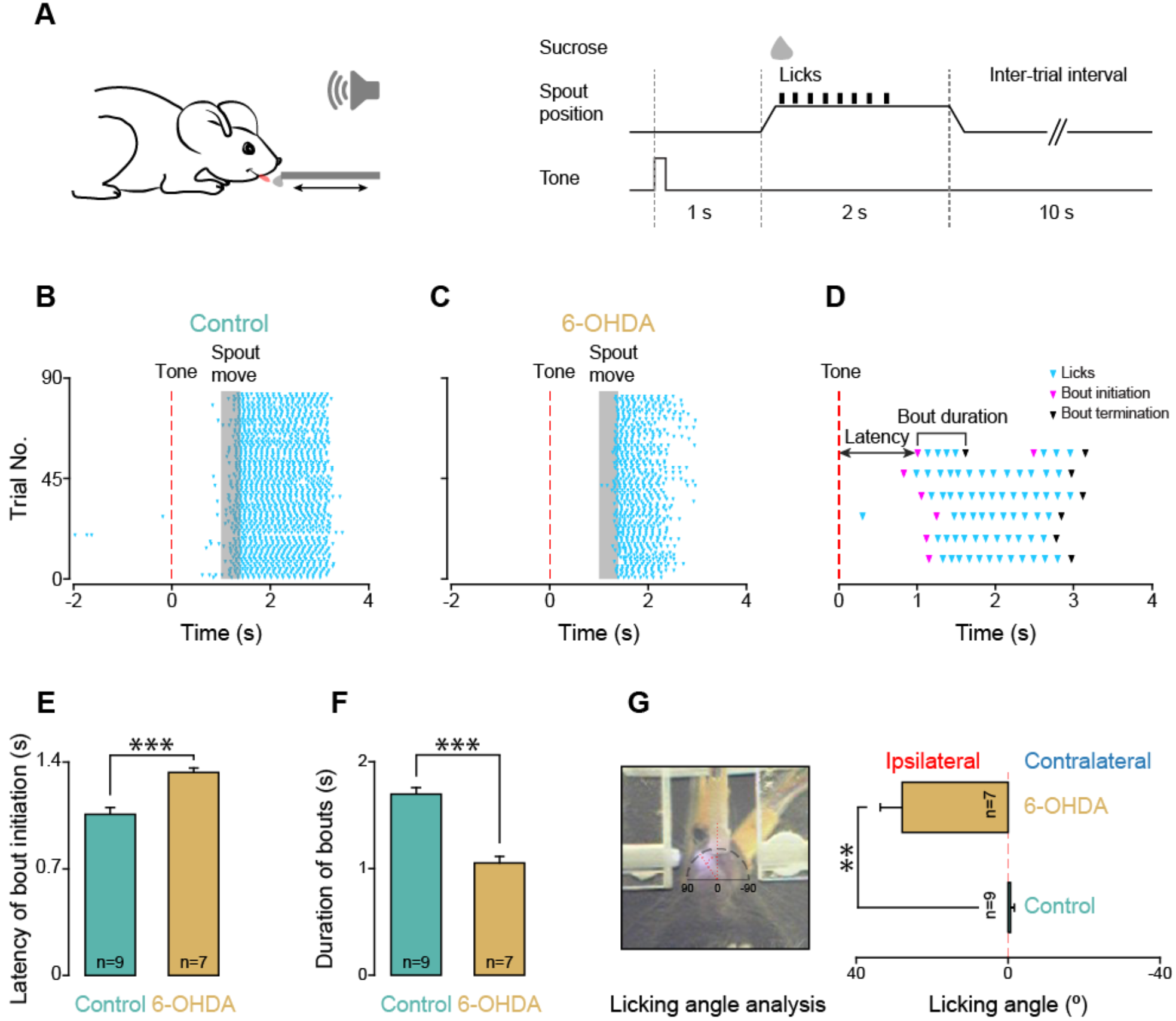
Licking deficits in 6-OHDA lesioned mice. **A**, *Left panel:* sketch showing a head-fixed mouse licking a spout to obtain sucrose. *Right panel:* schematic diagram of the experimental design for each trial. **B and C**, Representative raster plots of licking recorded from a control mouse (**B**) and a unilateral 6-OHDA lesioned (**C**) mouse performing the cued-licking paradigm. Dashed red vertical lines (time 0) indicate the onset of the auditory cue. Cyan triangles represent each individual lick. The gray shaded area highlights the movement of the spout. **D**, Representative raster plot of licking demonstrating bout analysis. A licking bout is defined as a train of at least three consecutive licks with an inter-lick interval shorter than 500 ms. Latency of bout initiation is defined as the latency of the first lick of a licking bout after tone onset. Cyan triangles represent each individual lick. Magenta triangles highlight the first lick of a licking bout (bout initiation) and black triangles highlight the last lick of a licking bout. **E and F**, Average values of latency of bout initiation (**E**) and duration of licking bouts (**F**) in control (n = 9 mice, blue) and 6-OHDA lesioned (n = 7 mice, brown) mice (**E and F**, t-test, *\*\*\*p* < 0.001). Error bars represent SEM. **G**, *Left panel,* a presentative snapshot showing a 6-OHDA lesioned mouse extending the tongue towards the licking spout. Note that the tongue protrudes on the right compared to the midline of the chin; *Right panel:* average values of the angles of tongue protrusion during licking in control (n = 9 mice, blue) and 6-OHDA lesioned (n = 7 mice, brown) mice (Welch’s corrected t-test, **p <0.01). Error bars represent SEM.

Altogether, these results demonstrate that mice with unilateral dopamine depletion have a longer latency to initiate a lick, a shorter duration of licking bouts, and a directional bias of the tongue toward the side ipsilateral to the lesion.

### Changes in cue responses and preparatory activity in ALM after dopamine depletion

Evidence from the literature points at the anterior lateral motor cortex (ALM) as the area responsible for modulating licking and controlling licking direction (Komiyama et al., 2010; Guo et al., 2014; Li et al., 2015; Li et al., 2016; Chen et al., 2017; Inagaki et al., 2018). To assess possible deficits in neural activity associated with dopamine depletion, we bilaterally recorded single units from ALMs of control (175 single units; n = 9 mice) and 6-OHDA lesioned mice (161 single units; n = 7 mice) engaged in the cued-licking paradigm described above. Units recorded from both hemispheres of control mice were pooled together. Units from 6-OHDA lesioned mice were analyzed separately depending on whether they were recorded on the side ipsilateral or contralateral to the site of the 6-OHDA lesion. We focused on firing rate modulations occurring in the interval from the onset of the cue to the initiation of licking bouts. We aligned neural activity either to the cue or to the bout initiation, and categorized neurons as cue responsive and/or preparatory depending on whether their firing changed shortly after the cue and/or just before licking (see methods). **Figure 3A** shows raster plots and PSTHs for two representative cue-responsive neurons from control mice: one excited and one suppressed by the auditory cue. We found that 41.7% of neurons (73 of 175 units) from control mice changed their firing rates within 500 ms from the onset of the cue. Differently, only 14.3% of neurons (12 of 84 units) from the ipsilateral side, and 19.5% of neurons (15 of 77 units) from the contralateral side, of 6-OHDA lesioned mice were cue responsive (**Figure 3B**). The differences in the proportion of cue responsive neurons among these three groups were significant (Pearson’s χ^2^ test, χ^2^(2) = 25.48, *p* = 2.6 × 10^−6^). Specifically, the proportion of cue-responsive neurons in the ipsilateral and contralateral side in 6-OHDA lesioned mice was similar (Pearson’s χ^2^ test, χ^2^(1) = 0.449, Bonferroni adjusted *p* = 1). However, it was significantly reduced from that observed in control mice (Pearson’s χ^2^ test, control vs ipsilateral, χ^2^(1) = 18.14, Bonferroni adjusted *p* = 6.2 × 10^−5^; control vs contralateral, χ^2^(1) = 10.67, Bonferroni adjusted *p* = 0.003).

**Figure 3.**
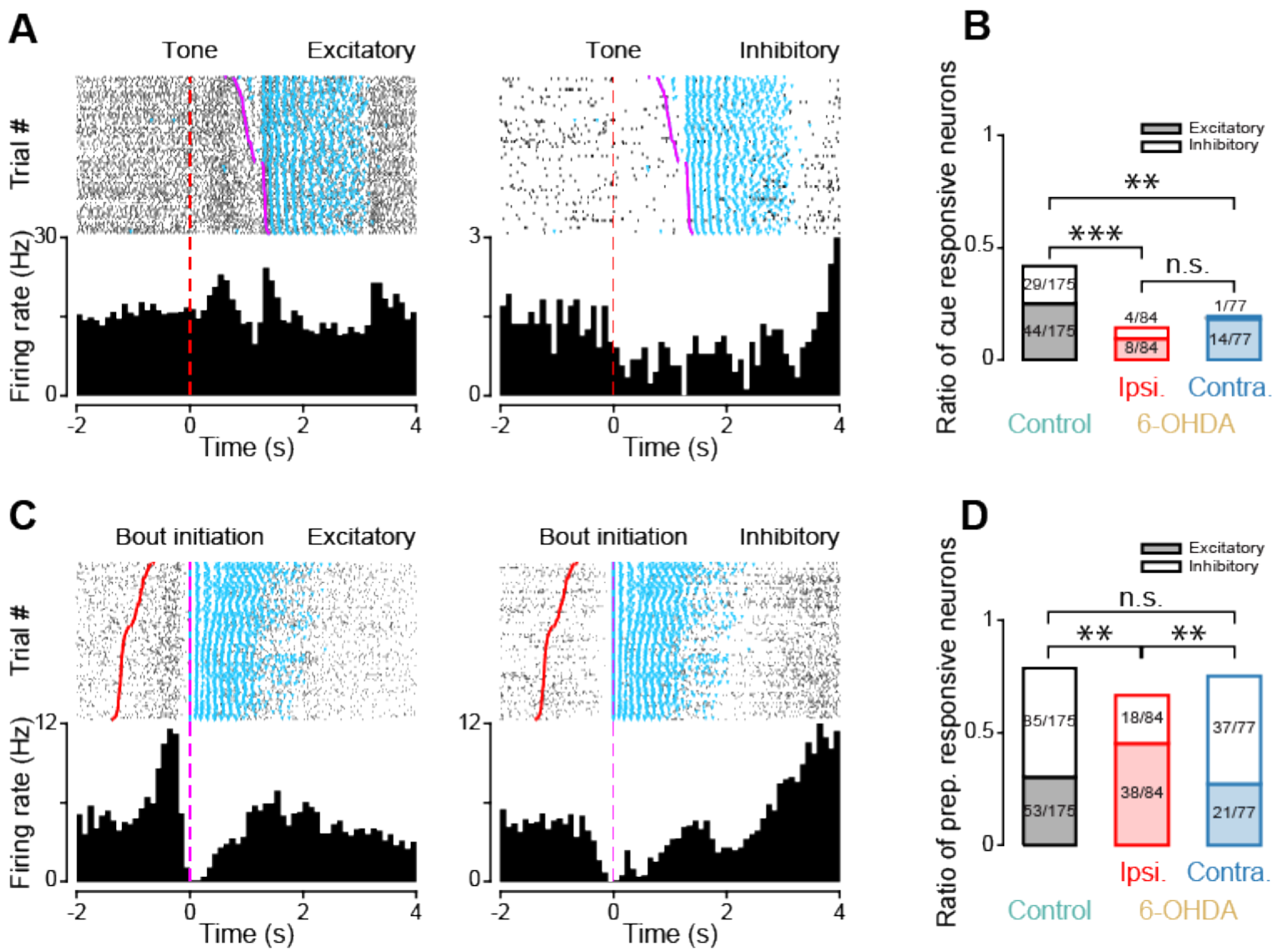
Cue responses and preparatory activity in ALM. **A**, Raster plots and PSTHs of neural activity recorded from two representative ALM neurons modulated by the cue within 500 ms from its onset. Dashed red vertical lines (time 0) indicate the onset of the auditory cue. Cyan markers represent each individual lick. Magenta markers represent the onset of each licking bout. Black ticks in raster plots represent individual action potentials. **B**, Proportion of cue responsive neurons in control mice (black) as well as ipsilateral (red) and contralateral (blue) sides of 6-OHDA lesioned mice *(post hoc* pairwise Pearson’s χ^2^ test for overall proportion of cue responsive neurons with Bonferroni correction, *** p<0.001, ** p<0.01, n.s. indicates not significant). **C**, Raster plots and PSTHs of neural activity recorded from two other ALM neurons modulated within 500 ms before licking bout initiation. Dashed magenta vertical lines (time 0) indicate the onset of the bout initiation. Cyan markers represent each individual lick. Red markers represent the onset of the cue. Black ticks in raster plots represent each action potential. **D**, Proportion of preparatory responsive neurons (filled: excitatory responses; empty: inhibitory responses) in control mice (black) as well as ipsilateral (red) and contralateral (blue) sides of 6-OHDA lesioned mice *(post hoc* pairwise Pearson’s χ^2^ test for excitation-inhibition ratio of preparatory responses with Bonferroni correction, ** p < 0.01, n.s. indicates not significant).

A large fraction of cue responsive neurons was also preparatory (90.4%, 66 of 73 units from control; 88.9%, 24 of 27 units from lesioned mice). However, not all preparatory neurons showed modulation of their activity by the onset of the cue: 52.2% (72 of 138) of the units from control and 78.9% (90 of 114) from lesioned animals did not show modulation by the cue. This difference indicates that, in a subset of neurons, preparatory activity started longer than 500 ms after the cue, thus closer to licking onset. **Figure 3C** shows raster plots and PSTHs of two representative neurons with preparatory activity recorded in control mice: the activity of one of the neurons is increased and that of the other neurons is suppressed before the initiation of a licking bout. In total, the percentage of neurons showing preparatory activity was 78.9% (138/175) in control, 66.7% (56/84) in the ipsilateral side and 75.3% (58/77) in the contralateral side of 6-OHDA lesioned mice (**Figure 3D**). Although the proportion of preparatory responses was similar across groups (Pearson’s χ^2^ test, χ^2^(2) = 4.50, *p* = 0.105), there were significant differences in the ratio of excitatory and inhibitory responses (Pearson’s χ^2^ test, χ^2^(2) = 16.06, *p* = 3.2 × 10^−4^). Specifically, neurons in the ipsilateral side of 6-OHDA lesioned mice showed a significantly larger proportion of excitatory responses when compared to neurons from control mice (ipsilateral side: 67.9% [38/56] excitatory, 32.1% [18/56] inhibitory; control: 38.4% [53/138] excitatory, 62.6% [85/138] inhibitory; Pearson’s χ^2^ test, χ^2^(1) = 17.72, Bonferroni adjusted *p* = 0.001), and from the contralateral side of 6-OHDA lesioned mice (ipsilateral side: see above; contralateral side: 36.2% [21/58] excitatory, 63.8% [37/58] inhibitory; Pearson’s χ^2^ test, χ^2^(1) = 10.20, Bonferroni adjusted *p* = 0.004).

Altogether, these results show that unilateral 6-OHDA lesions produce alterations in the proportion of cue responsive neurons and changes in the ratio of excitatory and inhibitory responses for preparatory activity.

### Slower onset of preparatory responses in ALM after dopamine depletion

Given the high prevalence of preparatory responses in our experimental conditions, we further analyzed them to extract possible differences in their time course. Since preparatory activity in ALM is important for planning tongue-related movements (Guo et al., 2014; Li et al., 2015; Inagaki et al., 2018), it is reasonable to expect that the slow onset of licking observed with dopamine depletion may relate to changes in the latency of preparatory activity. **Figure 4A** and **4B** show raster plots and PSTHs of four representative neurons with preparatory responses aligned to the onset of the cue: two from control mice (**Figure 4A**, left: excitatory, right: inhibitory) and two from ipsilateral side of 6-OHDA lesioned mice (**Figure 4B**, left: excitatory, right: inhibitory). **Figure 4C** and **4D** display the normalized responses (auROC, see methods) for all the preparatory neurons recorded from both hemispheres of control and lesioned mice. Visual inspection of the population activity suggests that the onset of preparatory firing may be delayed in 6-OHDA lesioned mice. This suggestion is corroborated by population PSTHs shown in **Figure 4E**. The latency of preparatory activity was directly quantified using a change point (CP) analysis approach (see methods). Response latency differed across conditions (Kruskal-Wallis Test, H_(2)_ = 30.68, p = 2.2 × 10^−7^). While neurons in the ipsilateral and contralateral side of 6-OHDA lesioned mice showed preparatory responses with comparable latencies (0.82 ± 0.06 s vs 0.70 ± 0.05 s, n = 56 and 58 respectively, *post hoc* Tukey HSD test, *p* = 0.184); the latency in both groups was longer than in control mice (ipsilateral side vs control: 0.82 ± 0.06 s vs 0.46 ± 0.03 s, n = 56 and 131 respectively, *post hoc* Tukey HSD test, *p* = 4.2 × 10^−7^; contralateral side vs control, 0.70 ± 0.05 s vs 0.46 ± 0.03 s, n = 58 and 131 respectively, *post hoc* Tukey HSD test, *p* = 0.003) (**Figure 4F** and **4G**).

**Figure 4.**
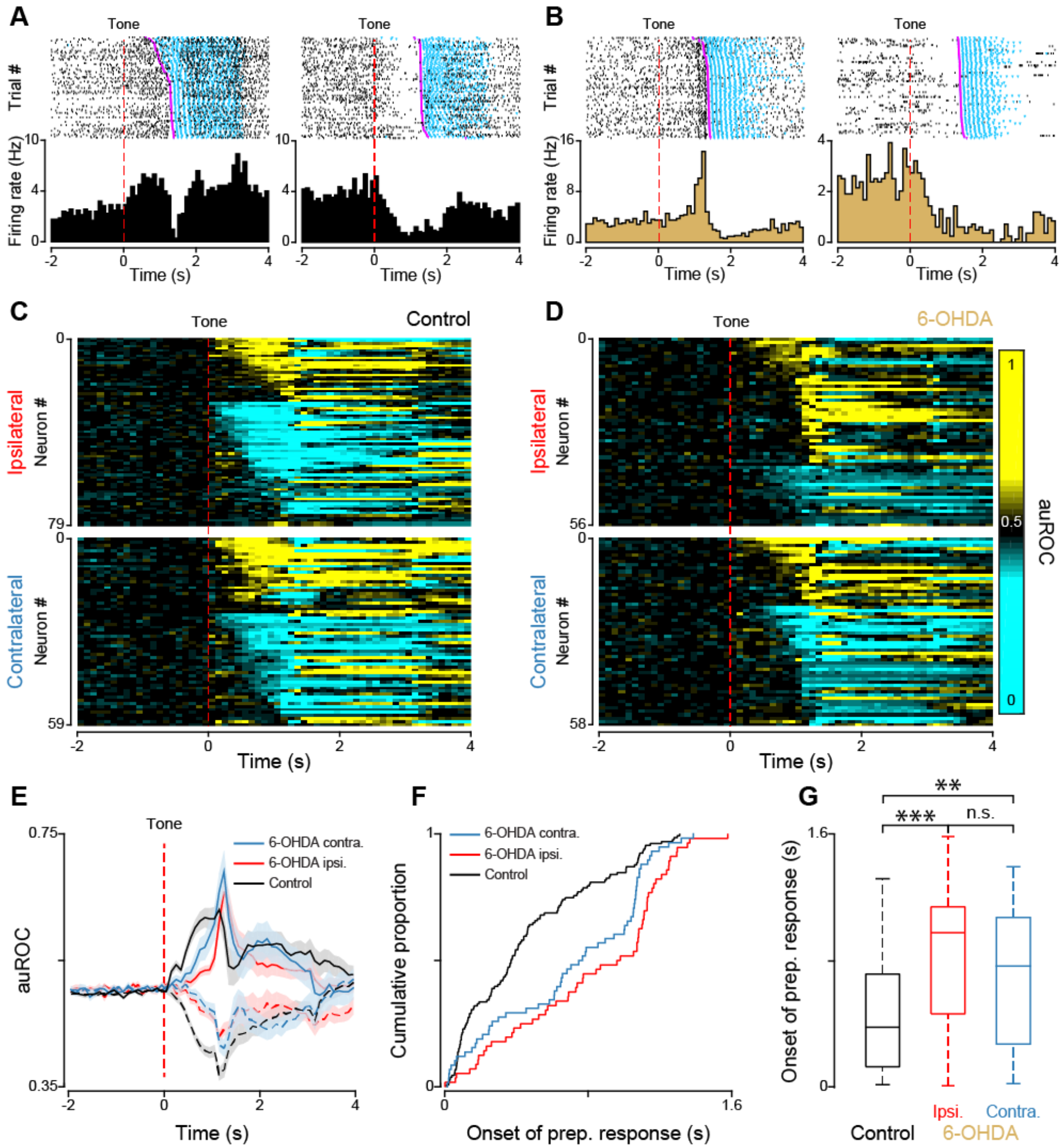
Timing of preparatory activity relative to the onset of the cue in control and 6-OHDA lesioned mice. **A and B**, Raster plots and PSTHs of neural activity recorded from four ALM neurons showing representative excitatory and inhibitory preparatory activity recorded from control (**A**) and 6-OHDA lesioned mice (**B**). Dashed red vertical lines (time 0) indicate the onset of the auditory cue. Cyan markers represent each individual lick. Magenta markers represent the onset of each licking bout. Black vertical ticks in raster plots represent action potentials. **C and D**, Population plots of all ALM neurons recorded from ipsilateral and contralateral sides in control (**C**) and 6-OHDA lesioned (**D**) mice. Each row represents a neuron and the color of each square along the x axis represents the normalized (auROC) firing rate within each 100 ms bin. Dashed red vertical lines (time 0) indicate the onset of the auditory cue. **E**, Population PSTHs of excitatory and inhibitory preparatory responses from control mice (black; data from ipsilateral and contralateral ALM were pulled together), ipsilateral (red) and contralateral (blue) sides of 6-OHDA lesioned mice. The dashed red vertical line (time 0) indicates the onset of the auditory cue. The shadow area around each curve represents the corresponding SEM. **F and G**, Cumulative distributions (**F**) and boxplots (**G**) for the latency of preparatory activity relative to the cue onset in control (black), ipsilateral (red) and contralateral (blue) sides of 6-OHDA lesioned mice (Kruskal-Wallis test, post hoc Tukey HSD test, ** p<0.01, *** p<0.001, n.s. indicates not significant).

To investigate preparatory activity relative to the onset of movement, we re-aligned spikes to the initiation of a licking bout (**Figure 5A** and **5B**). Visual inspection of population PSTHs suggests a possible difference in the latency of preparatory activity relative to licking initiation (**Figure 5C**). Indeed, CP analysis revealed significant differences across conditions (Kruskal-Wallis test, H (2) = 12.33, *p* = 0.002) (**Figure 5D** and **5E**). There were no significant differences in the onset of preparatory activity relative to the initiation of licking between neurons in control mice and in the contralateral side of 6-OHDA lesioned mice (−0.73 ± 0.04 s vs −0.75 ± 0.07 s, n = 131 and 56 respectively, *post hoc* Tukey HSD test, *p* = 1). However, the onset of preparatory activity in neuron from ipsilateral ALM in 6-OHDA lesioned mice was significantly closer to the initiation of licking when compared to that in control mice (−0.51 ± 0.06 s vs −0.73 ± 0.04 s, n = 54 and 131 respectively, *post hoc* Tukey HSD test, *p* = 0.002), and contralateral ALM of 6-OHDA lesioned mice (−0.51 ± 0.06 s vs −0.75 ± 0.07 s, n = 54 and 56 respectively, *post hoc* Tukey HSD test, *p* = 0.01).

**Figure 5.**
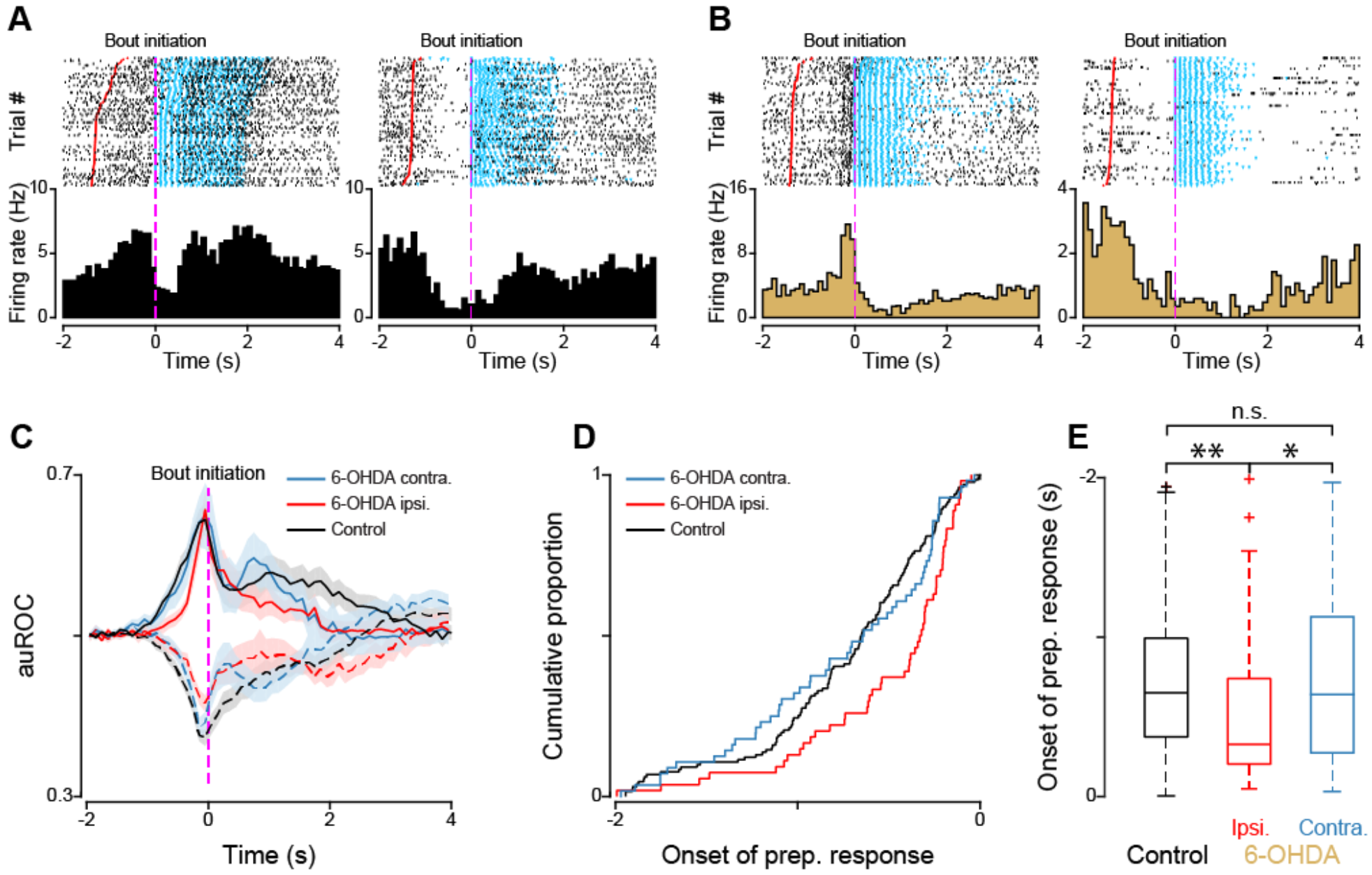
Timing of preparatory activity relative to the onset of a licking bout in control and 6-OHDA lesioned mice. **A and B**, Raster plots and PSTHs of the same ALM neurons shown in **Fig. 4A** and **4B**, but re-aligned to licking bout initiation. Dashed magenta vertical lines (time 0) indicate the bout initiation, red markers indicate the onset of the auditory cue, cyan markers represent each individual lick. Black ticks in the raster plots represent individual action potential. **C**, Population PSTHs of excitatory and inhibitory preparatory responses recorded from ALM neurons of control mice (black), ipsilateral (red) and contralateral (blue) sides of 6-OHDA lesioned mice. The dashed magenta vertical line (time 0) indicates the initiation of licking bouts. The shadow area around each curve represents the corresponding SEM. **D and E**, Cumulative distributions (**D**) and boxplots (**E**) for the latency of preparatory activity relative to the bout initiation in control (black), ipsilateral (red) and contralateral (blue) sides of 6-OHDA lesioned mice (Kruskal-Wallis test, post hoc Tukey HSD test, * p < 0.05, ** p<0.01, n.s. indicates not significant).

Altogether, neural recordings in 6-OHDA lesioned mice show that unilateral dopamine depletion induces changes in cue responsiveness and preparatory activity. There are fewer cue responsive neurons in lesioned animals. While the incidence of preparatory neurons was not affected, 6-OHDA lesions altered the balance between excitation/inhibition and delayed the timing of preparatory activity.

### D1 but not D2 receptor antagonism in ALM slows licking initiation

The results described above demonstrate significant alterations of neural activity in ALM following unilateral 6-OHDA lesions in the MFB. Are these changes epiphenomenal or indicative of a contribution of ALM to the licking deficits observed in 6-OHDA lesioned mice? To determine the link between dopaminergic modulation in ALM and licking deficits, we unilaterally and acutely infused D1 or D2 receptor antagonists into ALM of a new cohort of unlesioned mice (naïve) trained to perform the cued-licking paradigm. Infusion of a D1 receptor antagonist (SCH23390 hydrochloride, 5 μg/μl) significantly increased the latency of bout initiation (1.18 ± 0.05 s vs 1.47 ± 0.03 s, *n* = 7, paired t-test, *t*_(6)_ =-6.64, *p* = 5.6 × 10^−4^) (**Figure 6A**) and reduced the duration of licking bouts (1.85 ± 0.09 s vs 1.09 ± 0.13 s, *n* = 7, paired t-test, *t*_(6)_ = 9.62, *p* = 7.2 × 10^−5^) when compared to control, saline-infused, mice (**Figure 6B**). The licking angle, however, was not significantly affected (SCH23390 vs saline: 3.6 ± 0.8 deg vs 1.0 ± 1.1 deg, *n* = 7, paired t-test, *t*_(6)_ = 1.61, *p* = 0.158) (**Figure 6C**). Differently, ALM infusion of a D2 antagonist (raclopride tartrate salt, 5 μg/μl) did not significantly affect the latency of bout initiation (raclopride vs saline: 1.16 ± 0.05 s vs 1.15 ± 0.04 s, *n* = 9, paired t-test, t_(8)_ = 0.20, *p* = 0.85) (**Figure 6D**), licking bout duration (raclopride vs saline: 1.66 ± 0.11 s vs 1.65 ± 0.08 s, *n* = 9, paired t-test, t_(8)_ = 0.09, *p* = 0.93) (**Figure 6E**) or licking angle (raclopride vs saline: 0.1 ± 0.8 deg vs 0.5 ± 1.1 deg, *n* = 6, paired t-test, t_(8)_ = 0.55, *p* = 0.60) (**Figure 6F**).

**Figure 6.**
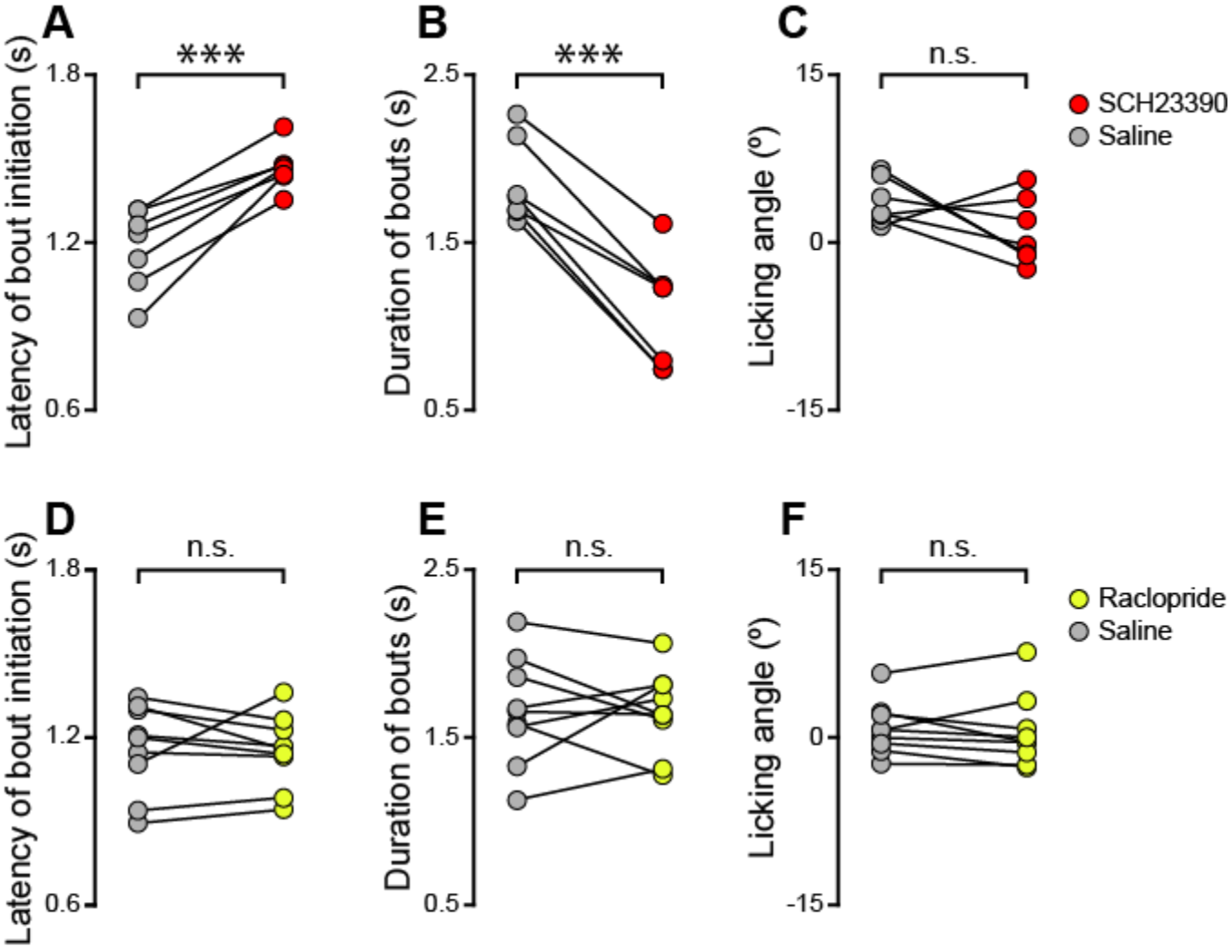
Effects of acute, local infusions of D1 and D2 receptor antagonists in ALM on licking. A, B and C, Latency of bout initiation (**A**), duration of licking bouts (**B**) and licking angle (**C**) recorded after unilateral infusion in ALM of saline (gray circles) or the D1 receptor antagonist, SCH23390 (red circles) (n = 7, paired t-test, *** p<0.001, n.s. indicates not significant). D, E and F, Latency of bout initiation (**D**), duration of bouts (**E**) and licking angle (**F**) recorded after unilateral infusion in ALM of saline (gray circles) or the D2 antagonist, raclopride (yellow circles) (n = 9, paired t-test, n.s. indicates not significant).

These data demonstrate that acute, unilateral blockade of D1, but not D2, dopaminergic signaling in ALM of naïve mice reproduces the behavioral impairments in licking initiation and duration observed in 6-OHDA lesioned mice, but not the ipsilateral bias in licking direction.

### Blockade of dopamine D1 receptor in ALM affects cue responses and preparatory activity

To identify the neural correlates of licking deficits observed after acute, local D1 receptor blockade, we infused SCH23390 (or saline) unilaterally into ALM of mice performing the cued-licking paradigm, and recorded single unit activity from the same side of the cortex. Unilateral infusion of D1 receptor antagonist significantly reduced the proportion of cue responsive neurons compared to saline infusions (SCH23390: 11.4% [5/44]; saline: 34.3% [12/35]; Pearson’s χ^2^ test, χ^2^(1) = 4.78, *p* = 0.029) (**Figure 7A**). Infusion of SCH23390 did not change the overall prevalence of neurons with preparatory activity (SCH23390: 72.7% [32/44], saline: 65.7% [23/35], Pearson’s χ^2^ test, proportion: χ^2^(1) = 0.182, *p* = 0.669), nor the relative proportion of excitatory and inhibitory response compared to control (SCH23390: 56.2% [18/32] excitatory, 43.8% [14/32] inhibitory; saline: 47.8% [11/23] excitatory, 52.2% [12/23] inhibitory; Pearson’s χ^2^ test, proportion: χ^2^(1) = 0.12, *p* = 0.731) (**Figure 7B** and **7C**). D1 receptors blockade did, however, affect the latency of preparatory activity, as suggested by visual inspection of population PSTHs (**Figure 7C, 7D** and **7G**). Quantification of the latency of preparatory activity relative to the cue revealed that D1 receptor antagonist infusion in ALM delayed its onset compared to control infusions (0.8 ± 0.09 s vs 0.44 ± 0.06 s, n = 21 and 31 respectively, Wilcoxon rank-sum test, W = 414, p = 0.008) (**Figure 7E** and **7F**). To compare the timing of preparatory activity relative to the onset of movement, we re-aligned spikes to the initiation of a licking bout. SCH23390 moved the onset of preparatory spiking closer to the initiation of licking compared to control (−0.52 ± 0.08 s vs −0.77 ± 0.10 s, n = 20 and 29 respectively, Wilcoxon rank-sum test, W = 403, P = 0.0496) (**Figure 7H** and **7I**).

**Figure 7.**
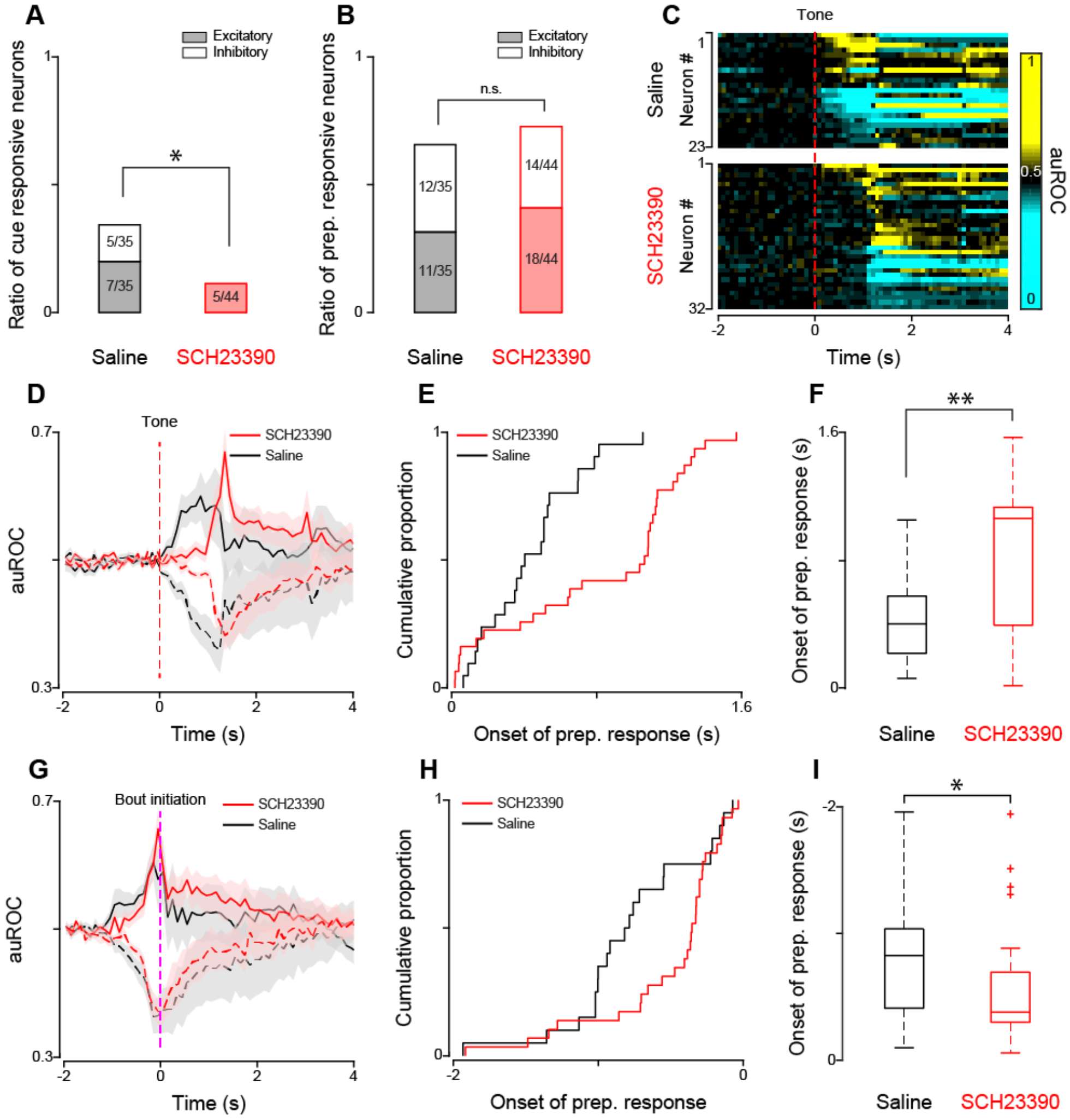
Effects of acute, local blockade of D1 receptor on patterns of single neuron activity in ALM. **A and B**, Proportion of neurons with cue responses (**A**) and preparatory responses (B) recorded with infusion of saline (black) and infusion of D1 receptor antagonist SCH23390 (red) in ALM (Pearson’s χ^2^ test, * p<0.05, n.s. indicates not significant). *C,* Population plot of preparatory activity in ALM recorded from mice with infusion of saline (top) and D1 receptor antagonist SCH23390 (bottom). Each row represents a neuron and the color of each square along the x axis represents the normalized (auROC) firing rate within each 100 ms bin. The dashed red vertical line (time 0) indicates the onset of the auditory cue. **D**, Population PSTH of preparatory activity after the infusion of saline (black) and SCH23390 (red). The dashed red vertical line (time 0) indicates the onset of the auditory cue. The shadow area around each curve represents the corresponding SEM. **E and F**, Cumulative distributions (**E**) and boxplots (**F**) for the latency of preparatory activity relative to the cue after the infusion of saline (black) and SCH23390 (red) (Wilcoxon rank-sum test, ** p<0.01). **G**, Population PSTH of preparatory activity with the infusion of saline (black) and SCH23390 (red). The dashed red vertical line (time 0) indicates the onset of licking bout initiation. The shadow area around each curve represents the corresponding SEM. **H and I**, Cumulative distributions (**H**) and boxplots (**I**) of the latency of preparatory activity relative to the licking bout initiation with the infusion of saline (black) and SCH23390 (red) (Wilcoxon rank-sum test, * p<0.05).

Altogether, these results show that acute intra-ALM infusion of a D1 receptor antagonist not only reproduces the slower licking initiation, but also recapitulates the reduction of cue responsive neurons and the slower onset of preparatory activity observed in 6-OHDA lesioned mice. Interestingly, neither lateral deviation of the tongue, nor changes in the proportion of excitatory and inhibitory responses were observed in animals infused with the antagonist.

## DISCUSSION

The results presented here provide behavioral, pharmacological and electrophysiological evidence showing a link between dopaminergic transmission in ALM, dysfunction of ALM neural activity and licking deficits. 6-OHDA lesioned mice trained to perform a cued-licking task showed delayed licking initiation, shorter duration of licking bouts and deviated tongue protrusion compared to controls. Single unit recordings revealed that unilateral dopamine depletion affects neural activity in ALM in several ways. First, it reduces the number of neurons activated by an anticipatory cue. Second, it changes the ratio between excitatory and inhibitory preparatory activity preceding movement, leading to more excitatory and fewer inhibitory modulations in the lesioned hemisphere of 6-OHDA lesioned mice. Finally, unilateral dopamine depletion results in delayed preparatory activity compared to controls. To determine whether disruption of cortical dopaminergic modulation directly caused licking deficits, we locally infused D1 or D2 receptor antagonists in ALM of unlesioned mice. Acutely antagonizing D1 receptors in ALM produced delayed licking initiation and shorter licking bouts. Single unit recordings after intra-ALM D1 blockade demonstrated that the behavioral deficits were associated with a reduction in the prevalence of cue responsive neurons and a delay in preparatory activity. Neither the lateral deviation of the tongue, nor the changes in the proportion of excitatory and inhibitory preparatory responses were reproduced by the infusion. Altogether, our data show a direct relationship between D1 receptor dopaminergic signaling in ALM, cue-evoked and preparatory firing and deficits in licking. More generally, these results suggest that cortical dopaminergic transmission may play a role in the genesis of some of the key symptoms of PD.

### Licking behavior after dopamine depletion

Patients with PD suffer from orolingual dysfunction, including tongue tremor and tongue weakness. Previous studies aimed at understanding how dopamine depletion affects tongue movement showed that unilateral 6-OHDA lesion of the MFB in rats significantly reduced tongue force and slightly increased the duration of pressing time during a tongue pressing test (Ciucci et al., 2011; Nuckolls et al., 2012). However, these experiments relied on a complex task in which rats were trained to press a disk with their tongue, and did not investigate natural licking or its latency of onset. Here, we studied tongue movements in the context of simple cued-licking paradigm. 6-OHDA lesioned mice displayed slower licking initiation, shorter bout duration (i.e. fewer licks per bout) and deviated tongue protrusion. The lesion did not affect inter-licking interval, demonstrating that the speed of each lick was not an issue in our animals.

These motor deficits could reflect an inability to control movement initiation, execution and termination. It is possible that some of the deficits observed in our task may be secondary to impairments in learning (Wise, 2004). For instance, delayed licking could derive from a reduced ability to associate a predictive cue with a reward. According to this view, longer latency to initiate movement would emerge from a weaker associative strength of the anticipatory signal or from the lack of anticipatory dopaminergic signaling. While we do not exclude underlying learning deficits in 6-OHDA lesioned mice, the results from acute unilateral infusions of D1 receptor antagonists in ALM emphasize the importance of real-time dopaminergic activity in the cortex in initiating movement.

ALM is known to regulate the direction of movement. Unilateral silencing of ALM activity, either with pharmacological or optogenetic approaches, causes deviation of the tongue on the ipsilateral side (Li et al., 2015). However, our local pharmacological manipulations demonstrate that this effect cannot be produced by acute, unilateral intra-ALM impairments in dopaminergic transmission. Hence, tongue deviation in 6-OHDA lesioned mice may result either from the effects of chronic unilateral disruption of ALM dopaminergic transmission, or by deficits that initiate in other nodes of the cortico-striatal loop (Von Voigtlander and Moore, 1973).

Altogether, our experiments establish active licking in mice as a model for studying motor deficits in the context of dopamine depletion, and point to the importance of D1 receptor signaling in ALM for mediating initiation and termination of tongue movements.

### Motor cortex and Parkinson’s Disease

Motor cortical activity is abnormal in PD patients and in animal models of PD (Lindenbach and Bishop, 2013). Changes in general excitability, excitation/inhibition balance, and timing have been described in the motor cortex during movement preparation or execution (Escola et al., 2003; Lindenbach and Bishop, 2013; Pasquereau et al., 2015). Our results on ALM fit with the existing literature and significantly extend it.

We showed that unilateral 6-OHDA lesion of the medial forebrain bundle impacts activity in the ALM. There was a significant reduction in the proportion of neurons whose firing rates showed modulation by the cue predicting the arrival of the spout. This result is consistent with the hypothesis of hypo-activation of motor cortex in PD and with recordings from MPTP-treated monkeys showing fewer cue responsive neurons in lesioned animals compared to controls (Escola et al., 2003). Although the total number of neurons changing their firing rates just before licking (i.e., preparatory neurons) was not affected by unilateral 6-OHDA lesion, we observed alterations in the ratio of excitatory and inhibitory modulations. Changes in excitation and inhibition were described in motor cortices of PD patients using paired-pulse transcranial magnetic stimulation (Lefaucheur, 2005; Lindenbach and Bishop, 2013). Our comparison of excitatory and inhibitory preparatory activity revealed a reduction in the proportion of neurons inhibited and an increase in the proportion of neurons excited prior to movement, a result consistent with the decrease of GABAergic tone observed in PD patient (Ridding et al., 1995). Finally, in addition to the changes described above, we observed deficits in the timing of preparatory activity. Preparatory activity in 6-OHDA lesioned mice had a longer latency from the cue compared to control mice, consistent with the delayed onset of the licking initiation observed after 6-OHDA lesion. Unilateral dopamine depletion affected the timing of preparatory activity also when spiking was aligned to the onset of licking. These changes in timing of neural activity were also observed in primate models of PD and in human PD patients (Doudet et al., 1990; Pasquereau et al., 2015). Specifically, in a reaction time task, PD patients showed a longer latency in initiating movement paralleled by a slower buildup of neuronal activation over the motor cortex (Dick et al., 1989; Mazzoni et al., 2012).

The results from acute D1 receptor blockade experiments provide very important information regarding the relationship between firing abnormalities in the cortex and licking deficits. They demonstrate that the reduction of cue responsive neurons and the delaying of preparatory activity in ALM can be sufficient to generate changes in motor systems leading to delayed licking. Furthermore, the lack of changes in balance between excitatory and inhibitory preparatory activity is evidence that this abnormality has limited causal role with regard to licking timing, and perhaps is more involved in tongue deviation (a symptom not present after local manipulations of ALM).

Altogether, our results show changes in ALM activity consistent with those described in PD patients and validate the study of ALM control of licking as a model for understanding the cortical involvement in PD.

### Dopaminergic modulation of cortical activity

Motor cortices receive direct dopaminergic innervation from the midbrain and dopaminergic inputs are known to play an important role in cortical plasticity and motor skill learning (Gaspar et al., 1991; Molina-Luna et al., 2009; Hosp et al., 2011; Guo et al., 2015). Dopamine exerts its function through five different receptors which are grouped into D1-like and D2-like receptors (Jaber et al., 1996). While both D1 and D2 receptors in motor cortex are important for modulating cortical plasticity and motor skill learning (Molina-Luna et al., 2009; Guo et al., 2015), here we show that dopaminergic signaling via D1, but not D2, receptors in ALM is required for modulating licking initiation and maintenance. This discrepancy may reflect the multiple functions of dopaminergic modulation in cortex. Our results indicate that acute D1 receptor signaling in ALM plays a role in modulating licking initiation and the timing of preparatory activity. This suggestion is consistent with recent findings showing transient activation of dopaminergic neurons before self-paced movement initiation (Jin and Costa, 2010; Howe and Dombeck, 2016; da Silva et al., 2018) and with experiments showing that optogenetic manipulation of transient dopaminergic activity can causally affect movement initiation (da Silva et al., 2018). In addition, our results on D1 receptor modulation of licking initiation dovetail nicely with existing literature in primates (Sawaguchi, 1995) and with data showing the importance of D1, but not D2, dopaminergic signaling in prefrontal cortex for the temporal control of action (Narayanan et al., 2012).

Our experiments clearly point at ALM D1 receptors as important in licking initiation and in modulating cue responses and preparatory firing in ALM. How activation of D1 receptors contribute to the patterns of activity observed in the ALM of mice performing a cued-licking paradigm remains to be seen and will be the subject of future investigations.

## Acknowledgements

The authors would like to acknowledge Dr. Arianna Maffei, Dr. Craig Evinger, Dr. Joshua L. Plotkin, past and present members of the Fontanini and Maffei’s laboratories for their feedback and insightful comments. This work has been supported by the Thomas Hartman Foundation for Parkinson Research.

## Notes

**Conflicts of interest:** The authors declare that no competing interests exist.

